# Ecological Role of the Heterotrophic Protist *Aurantiochytrium* (Labyrinthulomycetes) as a Key Consumer of Viral-Induced Dissolved Organic Matter Following the Lysis of the Red Tide-forming Microalga *Heterosigma akashiwo*

**DOI:** 10.64898/2026.04.06.716758

**Authors:** Shuhe Chen, Miho Aoki, Keishiro Sano, Keigo Yamamoto, Yoshitake Takao, Ryoma Kamikawa, Takashi Yoshida

## Abstract

Marine algal blooms play a vital role in oceanic carbon cycling, yet the ecological consequences of algal organic matter released following their collapse via viral infection are poorly understood. Recent studies have shown that viral infection dramatically alters the host’s intracellular metabolite composition, and the subsequent viral lysate selectively promotes the growth of specific prokaryotic populations. This study aimed to elucidate the effect of organic matter derived from healthy and virus-infected cells of the bloom-forming alga *Heterosigma akashiwo* on the growth of heterotrophic eukaryotes, specifically Labyrinthulomycetes. These marine protists are primarily saprotrophic or predatory and contribute to dissolved organic matter (DOM) decomposition and nutrient cycling. Our field monitoring in Osaka Bay over 12 months revealed that while the overall Labyrinthulomycetes community showed no clear seasonality, specific populations of the protists co-occurred with *Heterosigma akashiwo*. To mechanistically investigate the potential trophic linkage suggested by these field observations, a co-culture system comprising *H. akashiwo*, its specific virus (HaV53), and *Aurantiochytrium* sp. NBRC102614, used here as a model Labyrinthulomycete, was established. In the co-culture experiments, viral lysis of *H. akashiwo* led to a significant increase in the cell density of *Aurantiochytrium* sp., demonstrating that *Aurantiochytrium* can thrive on substrates derived from the virus-infected alga, such as viral-induced dissolved organic matter (vDOM). These findings highlight that heterotrophic Labyrinthulomycetes are one of key consumers of virus-modified organic matter, playing a pivotal role in carbon cycling following the collapse of harmful algal blooms and influencing carbon transfer in coastal microbial food webs.

**IMPORTANCE:** Marine ecosystems are tightly regulated by the interplay between microalgae, viruses, and heterotrophic eukaryotes, yet their roles within this network have long been underestimated. Accordingly, this study aimed to provide an overview of the dynamics of environmental microalgae and heterotrophic eukaryotes, namely *Heterosigma* species and Labyrinthulomycetes, and to elucidate the impact of virus-infected *Heterosigma akashiwo* on the growth and proliferation of *Aurantiochytrium* species within heterotrophic Labyrinthulomycetes. This study revealed the dynamics of several Labyrinthulomycetes species associated with *Heterosigma* populations in coastal marine environments and demonstrated that *Aurantiochytrium* species have the capacity to redistribute carbon, such as by utilizing vDOM released during the termination of *Heterosigma* blooms via viral infection, thereby repositioning *Aurantiochytrium* from a passive component of *Heterosigma* viral infection toward an active ecological agent that facilitates energy transfer and contributes to the maintenance of microalgal community dynamics. Overall, this work provides new insights into the fate of virus-infected *Heterosigma* in coastal marine systems mediated by heterotrophic Labyrinthulomycetes, particularly *Aurantiochytrium* species, thereby filling an important knowledge gap in microbial ecology.

## INTRODUCTION

The oceanic photosynthesis-mediated productivity driven by microalgae forms the foundation of the marine food chain via carbon fixation (1, 2). In addition, the dissolved organic matter (DOM) derived from the microalgae (3) is utilized by heterotrophic microorganisms, which in turn influence and shape of marine ecosystems by reintegrating these organic substances into the food web (4). However, several microalgal genera, such as *Gonyaulax* (5), *Pseudo-nitzschia* (6), *Chaetoceros* (7), *Prymnesium* (8), *Chattonella* (9), and *Heterosigma* (10), have been reported to form harmful algal blooms (HABs) that pose a threat to aquatic animals and humans, resulting in severe economic losses (11, 12).

Generally, HABs do not persist indefinitely. Several natural factors contribute to the decline and eventual collapse of harmful algal blooms, among which viral infection (13) and predation by heterotrophic eukaryotic microorganisms (14) are considered key factors that accelerate the termination process. Viruses influence carbon flow primarily through two mechanisms: the viral shunt and the viral shuttle, while their impact is tightly regulated by host specificity. The viral shunt redirects organic carbon derived from microalgae into the microbial loop through viral lysis (15), whereas the viral shuttle promotes vertical carbon export via sedimentation of virus-infected microalgal aggregates (16). Specifically, viruses can enhance virion production by manipulating host metabolism (17), resulting in significant changes in the composition of viral-induced dissolved organic matter (vDOM) released by infected cells (18) and a shift in metabolite composition (19). Viral host specificity is a crucial factor in regulating the population and density of microalgae. This massive release of vDOM from infected cells, termed ’virocells’ has been shown to significantly alter the composition of the surrounding coastal bacterial communities (20), suggesting a selective feeding mechanism based on the altered DOM profile (21). Blooms caused by *H. akashiwo* (22–24) are, at least in part, regulated by its specific double-stranded DNA virus, *Heterosigma akashiwo* virus (HaV). HaV was first isolated from Nomi Bay, Japan, in 1996 (25), and subsequently, other HaV strains, such as HaV53, were isolated from coastal environments (26). Successful adsorption to the host cell surface was found to be essential for viral infection, and adsorption kinetics mainly determined host susceptibility (27). HaV infection has also been shown to influence microalgal lysis rates and host strain diversity, with the algicidal activity and stability of HaV found to be affected by temperature (28). Consequently, HaV infection is likely to not only directly alters the cellular composition of bloom-forming of *H. akashiwo*, but also indirectly impacts primary productivity and biogeochemical cycles by reshaping the planktonic food web.

In addition to viruses, heterotrophic eukaryotic microorganisms, key components of the microbial loop, play a vital role in terminating microalgal blooms through their predation activities. For example, some heterotrophic dinoflagellates such as *Oxyrrhis marina* can efficiently prey on *H. akashiwo* (29) through phagocytosis (30). This phenomenon also occurs in Labyrinthulomycetes, which have been found to prey on microalgae via its ectoplasmic net system (31) and are involved in organic matter degradation through the production of extracellular enzymes (32). This suggests that heterotrophic eukaryotic microorganisms can regulate the microalgal population in a direct way, a process traditionally thought to contribute to the termination of algal bloom (33). However, whether Labyrinthulomycetes are capable of utilizing *H. akashiwo* specifically through predation remains unclear, although this group of heterotrophic eukaryotes has been shown to thrive in organic matter-rich waters (34). Most Labyrinthulomycetes are capable of degrading microalgae by secreting extracellular enzymes (35, 36), which can be linked to the potential utilization of marine microalgae. This process, along with other increasing evidence (37), is considered to accelerate the release of DOM, which facilitates the carbon cycle in marine systems.

To date, field observations suggest a link between Labyrinthulomycetes and phytoplankton dynamics (38), though the mechanisms governing their interaction, particularly following bloom-collapse events, are still not fully understood. An increase in AuRNAV (a virus infecting *Aurantiochytrium* species, a member of the Labyrinthulomycetes) was observed following the *H. akashiwo* bloom, suggesting a potential increase in the abundance of its hosts, *Aurantiochytrium* species (39). In turn, this observation raises the possibility that Labyrinthulomycetes may directly utilize the vast amounts of vDOM released during virus-induced lysis of microalgae such as *H. akashiwo*. Though the selective stimulation of prokaryotic communities by viral lysate has been elucidated (21), the response of heterotrophic eukaryotes to this altered vDOM input remains an open question. Therefore, this study integrates field observations with controlled laboratory experiments to address this gap in the response of heterotrophic eukaryotes to the altered algal-derived vDOM. In particular, the dynamics of Labyrinthulomycetes and *Heterosigma* communities in Osaka Bay were monitored to identify potential ecological links. Additionally, a tripartite model system involving *H. akashiwo*, its lytic virus, and *Aurantiochytrium* NBRC102614 which was isolated from the coastal area and is amenable to cultivation, was established to mechanistically test the hypothesis that Labyrinthulomycetes can thrive on vDOM derived from virus-infected *H. akashiwo*. This dual approach aims to elucidate the role of Labyrinthulomycetes in carbon cycling following the termination of harmful algal blooms, by clarifying the underlying ecological mechanisms.

## RESULTS AND DISCUSSION

### Monthly dynamics of eukaryotes with special reference to Labyrinthulomycetes in Osaka Bay

Eukaryotic microbial dynamics and environmental parameters were monitored at a coastal station in Osaka Bay (**Fig. 1**) monthly over a 12-month period. Surface water temperatures ranged from 9.77°C in January 2022 to 26.99°C in August 2022, with an average of 18.59 °C. Salinity varied between 31.86 and 33.54 psu, with an average of 32.59 psu. Concentrations of Silicte, SiO_2_, ranged from 1.57 to 16.41 μmol/L (**Fig. S1**).

**FIG 1.**
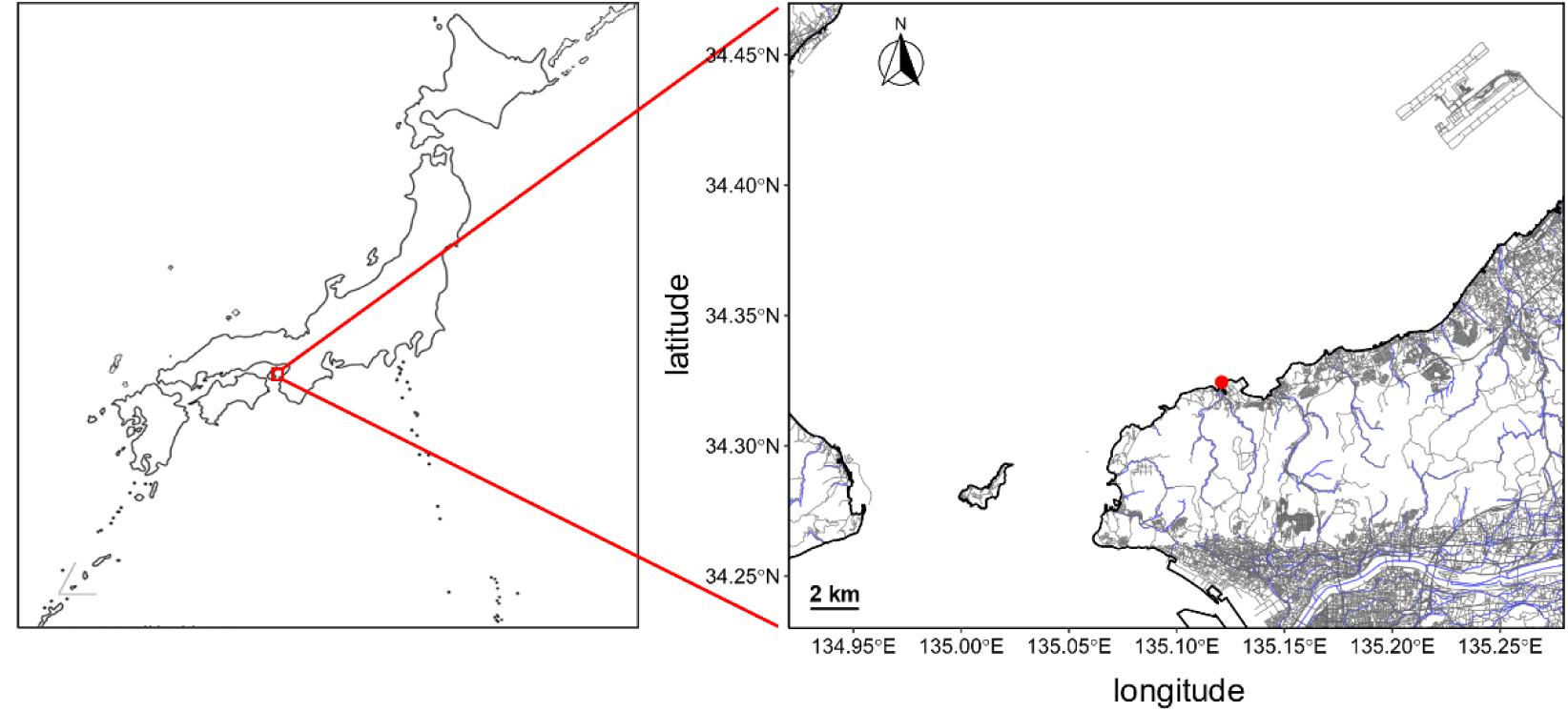
Sampling site. Located at the mouth of Osaka Bay (34°19’28”N, 135°7’15”E), pointed by red.

Taxonomic profiling of eukaryotic microbes at the rank 2 in the SILVA database based on 18S rRNA gene sequencing revealed that supergroup Stramenopiles (40.07% on average relative abundance) was the most dominant throughout the observation period, followed closely by Alveolata (37.54%). The high relative abundance of Alveolata was likely influenced by the elevated 18S rRNA gene copy numbers typical of dinoflagellates (40). Within Stramenopiles, a pronounced dominance of Ochyrophyta (85.66% on average), which is primarily composed of photoautotrophic species. Although the remaining 636 ASVs—representing the non-Ochrophyta stramenopiles ASVs—accounted for a mean relative abundance of 12.83%, they were predominantly heterotrophic. This contrast likely reflects differences in trophic strategies: autotrophs like Ochyrophyta act as primary producers, whereas heterotrophs rely on energy transfer through the food web, which is generally inefficient in coastal areas (5–27%) (41). A notable heterotrophic group within Stramenopiles was Labyrinthulomycetes, comprising 172 ASVs and accounting for 12.83% of the total relative abundance of Stramenopiles. In March 2022, a temporary increase in the relative abundance of Peronosporomycetes (29.76%) was observed. This group includes oomycete protists known to infect specific Bacillariophyceae species (42). This spike largely driven by a single ASV that contributed to 26.80% of the total Stramenopiles abundance, although it’s more detailed taxonomic identity remained unresolved.

Further investigation of Labyrinthulomycetes at the genus level indicated *Aplanochytrium* as the dominant genus (39.01% on average of total Labyrinthulomycetes), followed by *Oblongichytrium* (7.47% on average), *Thraustochytrium* (5.26% on average), and *Labyrinthula* (0.99% on average). Of the 172 Labyrinthulomycetes ASVs detected, only 69 (40.12% on average) were closely related to cultured strains listed in the SILVA v.138 database. The remaining 103 ASVs (59.88% on average) represented uncultured lineages, highlighting the limited ecological information available for this group. A total of 109 ASVs were identified as “dominant”, defined as >0.01% of the eukaryotic community at least one month. Among these, one *Aplanochytrium* ASV (ASV_L57) exhibited the highest average relative abundance (0.14%) and persisted across all months (**Fig. S2**). In contrast, most other dominant ASVs appeared transiently. Notably, the Labyrinthulomycetes community displayed no distinct seasonal patterns. This lack of clear seasonality may be attributed to their metabolic versatility; they secrete diverse extracellular enzymes capable of degrading broad spectra of organic matter, including terrestrial detritus and phytoplankton debris (36, 43), allowing them to maintain stable populations, independent of specific food source organisms. However, despite this lack of clear seasonal trend, Redundancy Analysis (RDA) indicated that the community structure was significantly influenced by temperature, whereas salinity and SiO_2_ had no significant effect (**Fig. S3**). This aligns with a previous finding (44), suggesting that while the group is present throughout the year due to substrate plasticity, their specific community composition or metabolic rates are thermally regulated.

Although Labyrinthulomycetes did not exhibit clear seasonal trends (**Fig. 2b**), co-occurrence analysis revealed potential biotic interactions. Specifically, eight Labyrinthulomycetes ASVs, including *Aplanochytrium* (ASV_L42), *Thraustochytrium* (ASV_L67, ASV_L72), and several uncultured species (ASV_L105, ASV_L119, ASV_L142, ASV_L170, and ASV_L171) showed strong positive correlations (Pearson correlation >0.7) with the dynamics of *Heterosigma akashiwo* (**Table S6**; **Fig. 4**). This suggests a potential ecological linkage, raising the question: can Labyrinthulomycetes utilize organic matter released from *Heterosigma* blooms? To mechanistically investigate this, we selected the Labyrinthulomycotes strain *Aurantiochytrium* sp. NBRC102614, as a model organism. This strain, isolated from a coastal region, is easily cultivable and serves as a suitable representative for studying host-virus-predator interactions.

**FIG 2.**
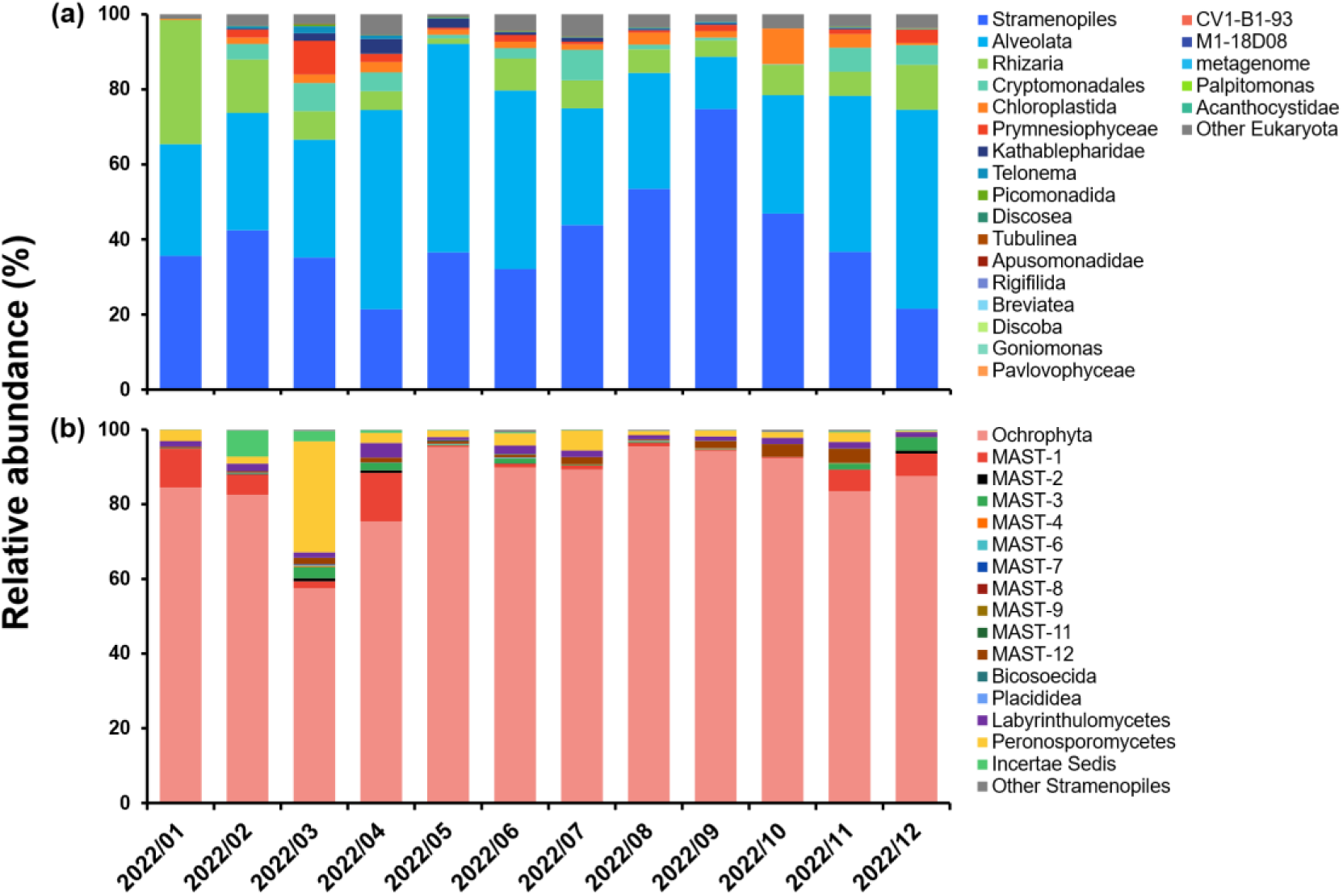
Relative abundance of eukaryotic lineages. (**a**) ASVs are classified at the supergroup/clade level. Other Eukaryota comprises those classified only to the domain level; (**b**) ASVs classified into the supergroup Stramenopiles were extracted, and the relative abundance of each ASV when classified at the class level was shown. Other Stramenopiles includes those classified only to the phylum level.

**FIG 3.**
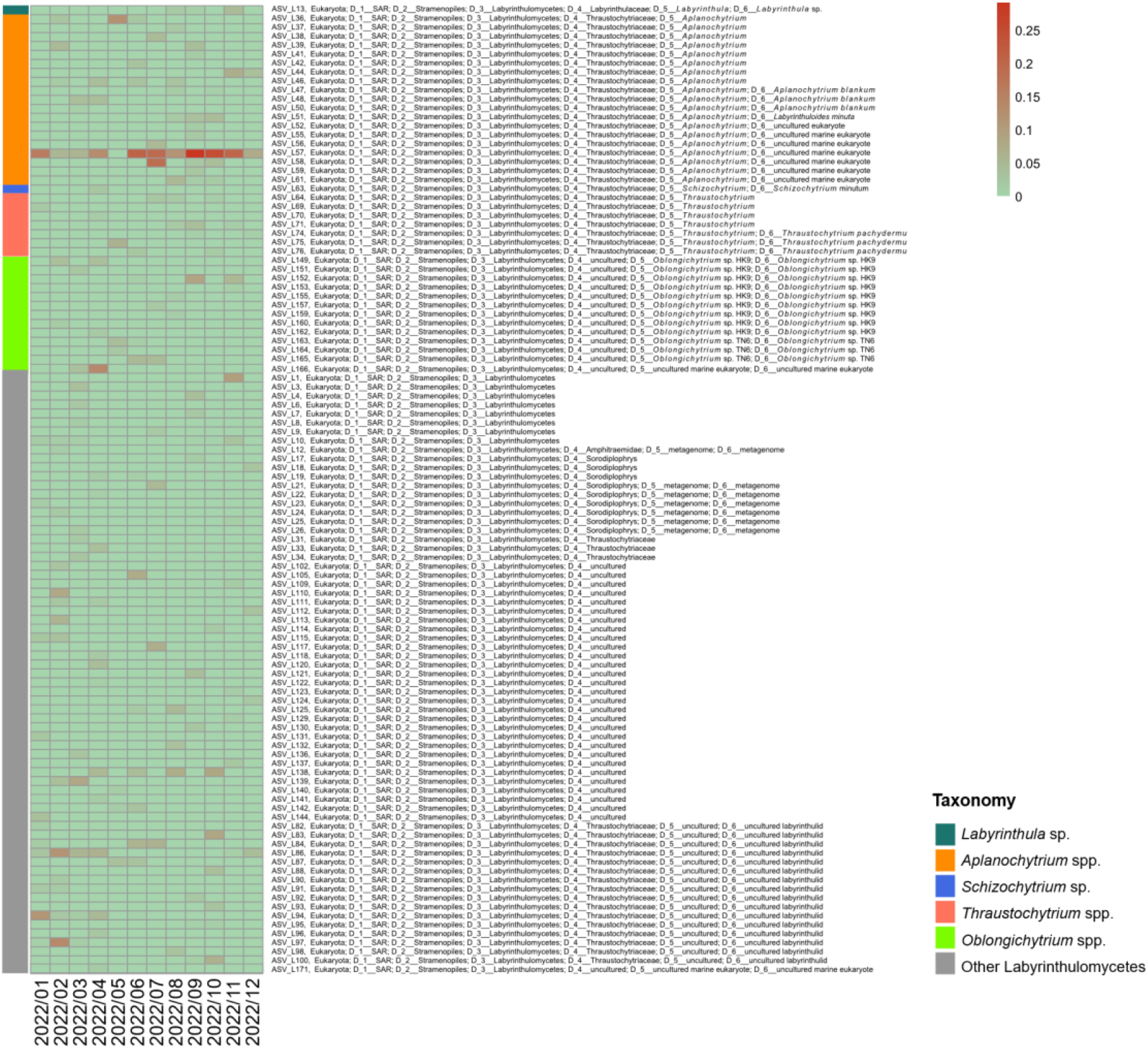
Relative abundance of 109 Labyrinthulomycetes-dominant ASVs. The color bar indicates relative abundance (%).

**FIG 4.**
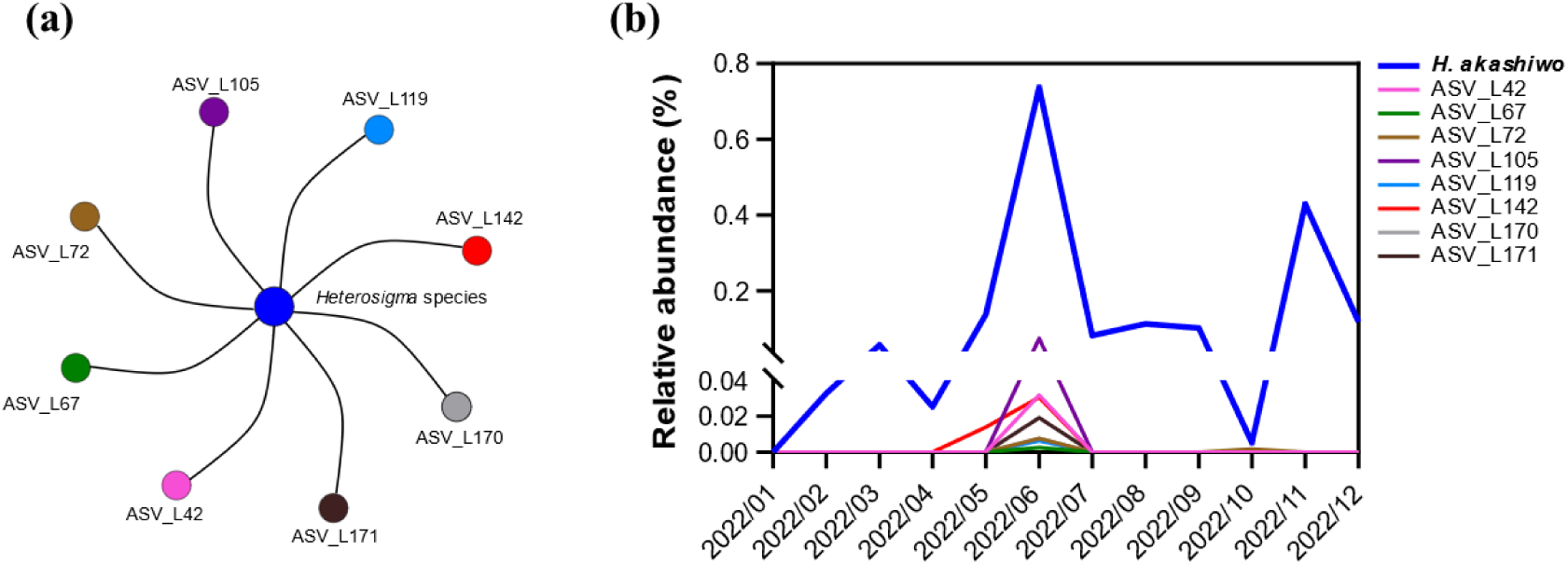
Co-occurring network between *H.* akashiwo and Labyrinthulomycetes species (**a**), and their relative abundances (**b**).

### Growth characteristics of *H. akashiwo* H93616 and *Aurantiochytrium* sp. NBRC102614, and validation of quantitative PCR

Growth monitoring with a flow cytometer (RF-500, Sysmex) analysis revealed that *H. akashiwo* reached a plateau by day 8, 4.46 × 10^4^ cells/mL (4.65 log_10_ cells/mL) on average reaching a maximum of 4.49 × 10^4^ cells/mL (4.65–4.66 log_10_ cells/mL) on day 12 (**Fig. S4a**). In contrast, in the co-culture, *Aurantiochytrium* sp. stabilized by day 5, 1.04 × 10^7^ cells/mL (7.01 log_10_ cells/mL) on average, reaching a maximum cell density of 1.28 × 10^7^ cells/mL on day 12 (7.11 log_10_ cells/mL) (**Fig. S4b**).

To quantify dynamics of *Aurantiochytrium* sp. NBRC102614, *H. akashiwo* H93616, and *H. akashiwo* virus HaV53 in the mixed culture, we developed qPCR assay targeting the β-tubulin gene for *H. akashiwo* and the *mcp* gene for HaV53. Standard curves showed strong linearity (R^2^ > 0.97) and acceptable amplification efficiencies (83.5 and 85.4%), confirming the reliability of the method for estimating cell and viral densities (**Fig. S5**).

### Proliferation of *Aurantiochytrium* stimulated by virus-induced lysis of *H. akashiwo*

In the triplet co-culture comprising *Aurantiochytrium* sp. NBRC102614, *H. akashiwo* H93616, and HaV53, the addition of HaV53 in viral-infected group (Infection) led to a rapid decline in algal cell density, with *H. akashiwo* falling below detection limits by day 3 (**Fig. 5b**). This confirms effective viral-mediated cell lysis given the non-infected control culture in which algal cell density remained stable at 1.69×10^5^ cells/mL on average (**Fig. 5b**), consistent with viral-mediated bloom collapse scenarios (47). Conversely, the growth of *Aurantiochytrium* was significantly stimulated in the viral-infected group, compared to the non-infected culture (Non-infection) and to the control (*Aurantiochytrium* species only, Control) (*p* < 0.05; **Fig. 5a**). Cell density of *Aurantiochytrium* in the infected culture peaked at 4.38×10^6^ cells/mL by day 6, suggesting that *Aurantiochytrium* responded to the vDOM released from lysed *H. akashiwo* cells.

**FIG 5.**
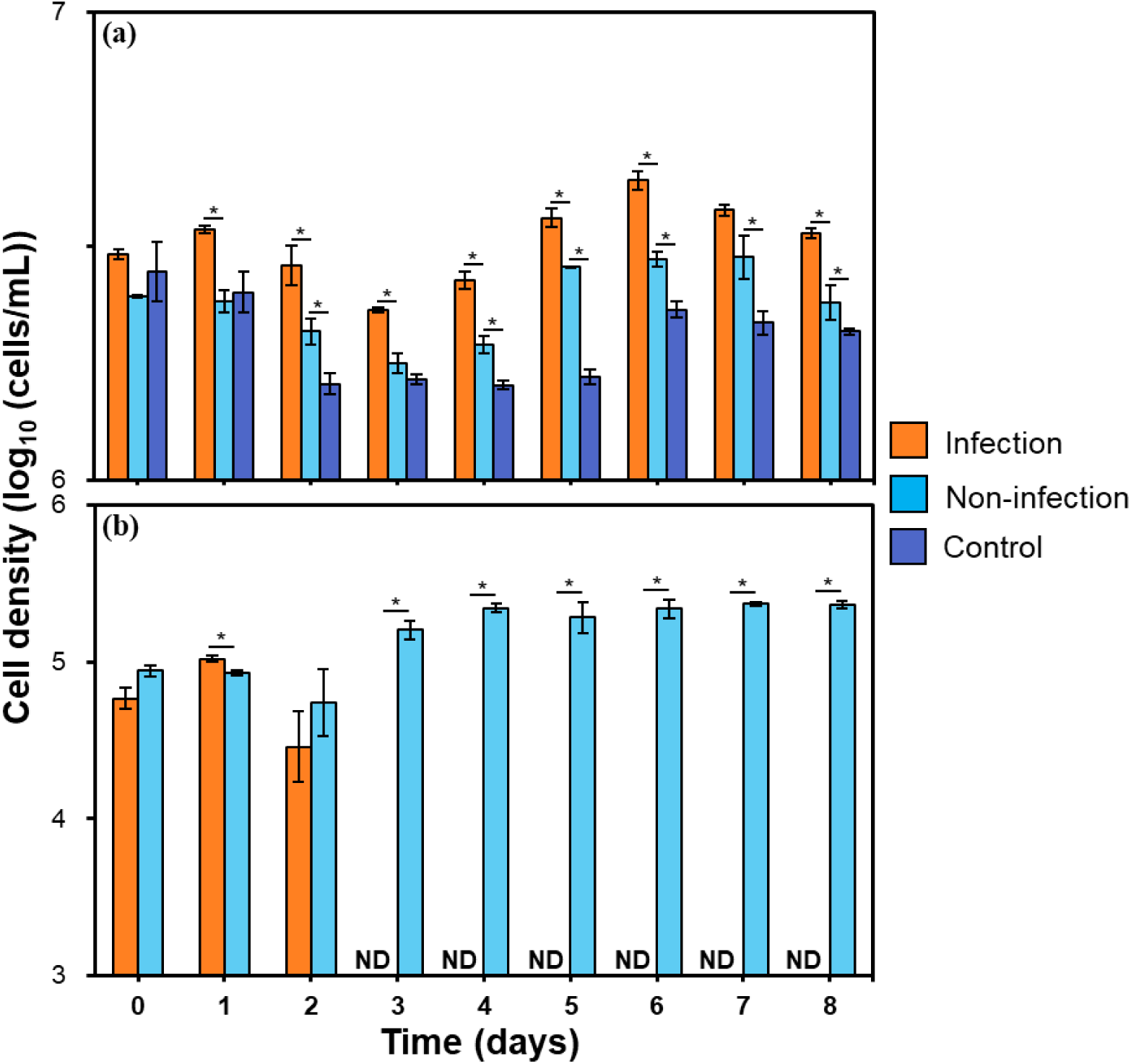
Changes in cell numbers during co-culture. (**a**) *Aurantiochytrium* sp. NBRC102614, and (**b**) *H. akashiwo* H93616. ‘**Infection’** refers to cultures of *Aurantiochytrium* sp. NBRC102614 + *H. akashiwo* H93616 + HaV53, while ‘**Non-infection**’ refers to cultures of *Aurantiochytrium* sp. NBRC102614 + *H. akashiwo* H93616, and ‘**Control**’ refers to cultures of *Aurantiochytrium* sp. NBRC102614 alone. The asterisk (_*_) at each time point indicates significant differences (*p* < 0.05) between treatments from the second day onward. Error bars represent Standard Error of the Mean (SEM), and ‘**ND**’ indicates values below the detection limit.

The significantly higher growth in the infected culture clearly indicates that viral lysis provides a more potent nutrient pulse than natural decay, considering the control culture where only natural senescence occurred. This supports the field observation based on environmental DNA sequencing that *Aurantiochytrium* signatures were absent from surface waters (**Fig. 3**) but may thrives on episodic organic inputs, such as algal lysis events (45).

### Quantification of *H. akashiwo* and its virus during viral infection

To confirm that the algal decline was due to lysis and not merely viral attachment, we monitored gene copy numbers of *H. akashiwo* and HaV53. The β-tubulin gene copy density of *H. akashiwo* decreased significantly from day 2 onwards in the virus-infected culture (Infection) (**Fig. 6a**), mirroring the flow cytometry data (**Fig. 5b)**. Concurrently, intracellular viral *mcp* gene copies were high on days 0 and 2, followed by an increase in extracellular *mcp* copies from day 2 (**Fig. 6b**), indicating viral release. However, the extracellular *mcp* gene copies plateaued after day 2. This absence of further accumulation suggests that viral particles were either not produced after host depletion or were rapidly degraded/adsorbed onto organic matter.

**FIG 6.**
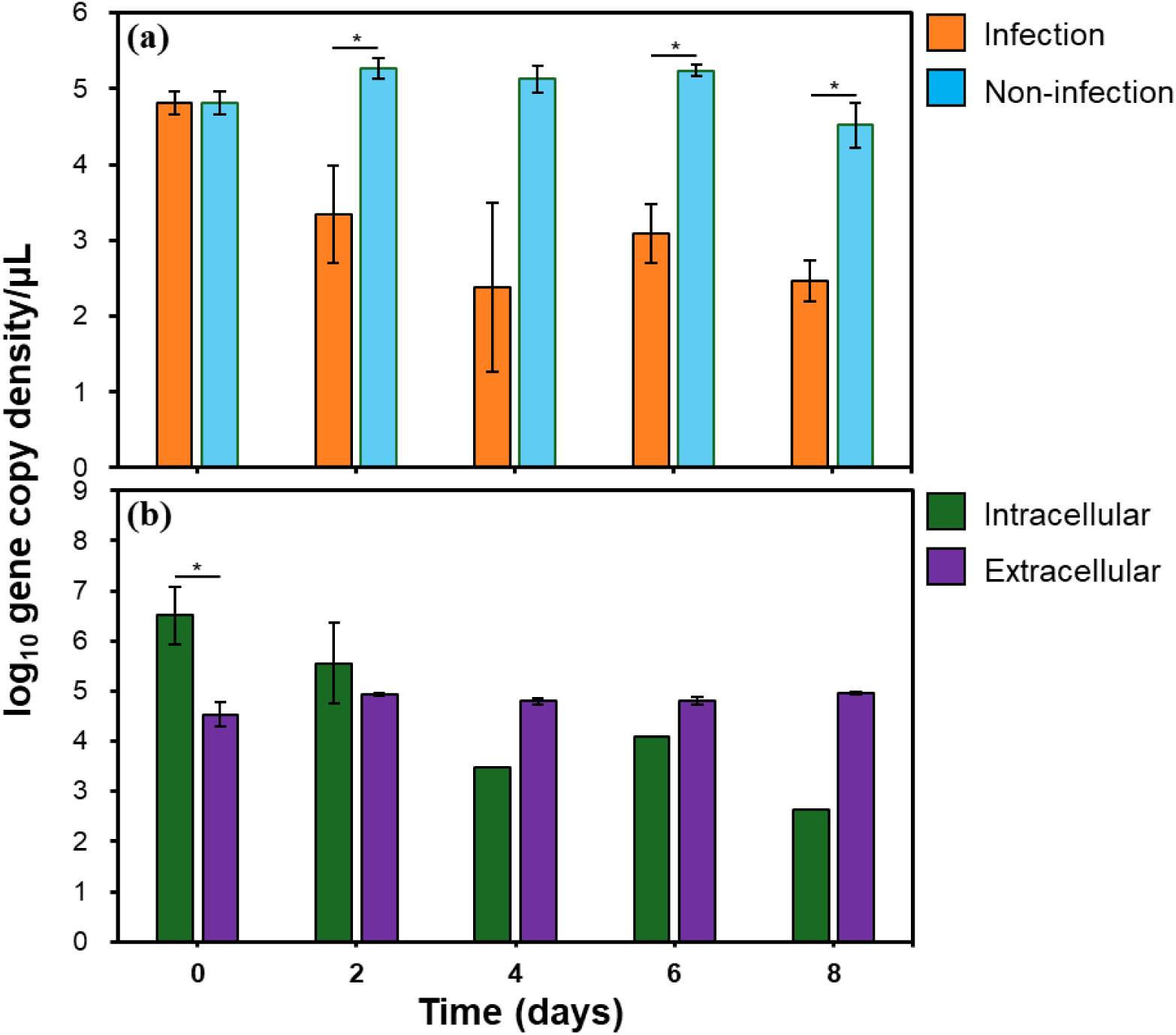
Changes in gene copy numbers during co-culture. (**a**) β-tubulin gene copy number of *H. akashiwo* H93616; (**b**) *mcp* gene of HaV53. **‘Infection’** refers to cultures of *Aurantiochytrium* sp. NBRC102614 + *H. akashiwo* H93616 + HaV53, whereas ‘**Non-infection’** refers to cultures of *Aurantiochytrium* sp. NBRC102614 + *H. akashiwo* H93616. **‘Intracellular’** refers to samples from the infection treatment collected by centrifugation, whereas **‘Extracellular’** refers to the supernatant collected from the same infection treatment. The asterisk (*) at each time point indicates significant differences (*p* < 0.05) between treatments; Error bars represent Standard Error of the Mean (SEM), and columns without error bars indicate that the number of data points is less than 2.

Collectively, these results support a mechanistic model in which viral lysis of bloom-forming algae releases vDOM and viral particles into the environment, stimulating heterotrophic protist activity in microbes, such as *Aurantiochytrium* sp. Such interactions not only influence microbial food web dynamics but may also accelerate the carbon flux by enhancing microbial decomposition and aggregation of algal detritus (46). Moreover, the observed rapid increase of *Aurantiochytrium* in parallel with *mcp* gene detection raises the possibility that this protist utilizes viral particles themselves as a direct nutrient source, which is consistent with emerging studies on viral shunt pathways and virus-derived nutrient recycling (47–49). These findings and possibilities highlight the underappreciated role of eukaryotic heterotrophs in post-lysis nutrient assimilation, suggesting that viruses not only terminate blooms but also restructure heterotrophic microbial community compositions and functions in marine ecosystems.

## MATERIALS AND METHODS

### Materials in this study

The red tide-forming microalga *H. akashiwo* strain H93616 and its virus HaV53 were isolated from Hiroshima Bay (26). Additionally, heterotrophic eukaryote *Aurantiochytrium* sp. strain NBRC102614 was originally isolated from Mikage Port in Kobe (50).

### Seawater sampling, total DNA extraction, rDNA amplicon sequencing, and assessment of environmental effects

Seawater samples were collected from seawater intake (34° 19’ 28”N, 135° 7’ 15”E) owned by the Fisheries Technology Center of the Osaka Prefectural Research Institute for Environment, Agriculture, and Fisheries, Osaka Prefecture (**Fig. 1**). The intake was located 5 m deep and 1.8 m above the seabed, enabling stable and continuous water collection throughout the year. Seawater was filtered using a membrane with a diameter of 142 mm and a pore size of 3.0 µm (EMD Millipore Isopore^TM^ Polycarbonate Membrane Filter, EMD Millipore, USA) to obtain the particle-associated (PA) fraction, yielding a total of 12 samples monthly from January to December 2022.

DNA was extracted following the protocol of the DNeasy PowerWater Kit (QIAGEN). Briefly, the filter was rolled into a cylinder with the cell-containing side facing inward and inserted into a 5 mL PowerWater DNA Bead Tube, which was then vortexed for 5 minutes to lyse the cells. Subsequently, the lysate was centrifuged at 500 g, and the supernatant was transferred to a new tube for centrifugation (13,000 g × 1 min) again, yielding a fresh supernatant that was transferred to a new tube, followed by the addition of 200 µL of Solution IRS. After gentle vortexing, the sample was incubated at 4 °C for 5 min and centrifuged at 13,000 g × 1 min at room temperature to yield the fresh supernatant. Supernatant was transferred to a new tube, followed by the addition of 650 µL of Solution PW3 for gentle vortexing. Subsequently, 650 µL of the supernatant was transferred to an MB Spin Column for centrifugation at 13,000 g × 1 min at room temperature, with the flow-through discarded and the process repeated until the entire supernatant was processed. The column was then washed with Solution PW4 and ethanol, finally eluted with 100 µL of EB Solution to yield the purified DNA. The concentration of the extracted DNA was measured using the Qubit^®^ dsDNA HS Assay (Life Technologies, Invitrogen division, Germany).

Amplicon sequencing of 18S rRNA genes was conducted from the PA fraction to analyze the eukaryotic community using a primer set based on the V8–V9 hypervariable region (51) with overhang adapter sequences added to each 5′ end (**Table S1**). Amplicons were sequenced using a MiSeq system with MiSeq V3 (2 × 300 bp) reagent kit (Illumina, San Diego, CA, USA). The adapter sequences were removed from the 18S rRNA gene amplicon sequences using Cutadapt in QIIME2 v.2022.11 (52). Denoising was performed using DADA2 package (53), to yield amplicon sequence variants (ASVs) (54). The taxonomy of the ASVs was assigned using a naïve Bayes classifier that was trained on the SILVA 138 database (SILVA 138 99% OTUs) using the regions of sequences that could be amplified using the abovementioned primer sets (54). ASVs resulting from 18S rRNA amplicon sequencing were filtered to remove singletons and Opisthokonta. Subsequently, ASVs were used to perform co-occurrence analysis using the Pearson correlation coefficient.

Additionally, the environmental parameters were also collected. The Redundancy analysis (RDA) (55) was performed to examine the environmental factors, namely temperature, and salinity, and SiO_2_ affecting the prokaryotic communities using the ggrepel v.0.9.6 and vegan v.2.6.8 packages in R v.4.4.2.

### Growth monitoring of *Heterosigma akashiwo* H93616 and *Aurantiochytrium* sp. NBRC102614

For growth monitoring of *H. akashiwo* H93616, 3 mL of *H. akashiwo* H93616 was transferred to 27 mL of sterilized f/2-Si medium using a 25 cm² VIOLAMO cell culture flask (ASONE) after pre-culture, and cultured in an artificial climate chamber at 20 °C under a 12 h light/12 h dark cycle with a light intensity of 100 µmol photons m^-^²s^-1^. The experiment was performed in triplicate. Subsequently, 960 µL of samples were collected from each flask daily from day 0 to day 17 and fixed with 40 µL of glutaraldehyde (final concentration, 1%). The fixed sample were subjected to a flow cytometer RF-500 (Sysmex) to determine cell density.

For *Aurantiochytrium* sp. strain NBRC102614, followed the method described in previous study, a colony was picked and inoculated into 20 mL of d-GPY liquid medium (containing glucose 2 g, peptone 1 g, and yeast extract 0.5 g in 1 L artificial seawater) following the method described in a previous study (56), and incubated for 2–3 days until reaching the logarithmic growth phase. Subsequently, 3 mL of the *Aurantiochytrium* sp. NBRC102614 culture was transferred to 27 mL of sterilized d-GPY medium and cultured in a growth chamber under controlled conditions (20 °C, 12-hour light /12-hour dark cycle, 100 µmol photon/m²/s). During cultivation, 10 µL samples were collected from each flask from day 0 to day 13 for cell density determination using a hemocytometer (WAKENYAKU) and a biological microscope (CX-43, OLYMPUS). All cell density data for *H. akashiwo* H93616 and *Aurantiochytrium* sp. NBRC 102614 were log-transformed.

### Establishment of a quantitative PCR for *H. akashiwo* and its virus HaV53

To establish a method for accurately quantifying the abundance of *H. akashiwo* and particularly its virus, HaV53, a quantitative PCR (qPCR) assay was developed in this study. Briefly, *H. akashiwo* H93616 was cultured for 3 days with an initial cell density of 10^4^ cells/mL. Subsequently, samples were collected for cell density determination using a hemocytometer (Wakenyaku) and a biological microscope (CX-43, OLYMPUS). DNA was then extracted by the xanthogenate-SDS method (57) with XS buffer (**Table S3**). For HaV53, viral DNA was extracted from collected samples (centrifugation at 20,000 g × 10 min) using the same method as described above (57). All extracted DNA was dissolved in the virus-free water prepared by filtering ultrapure water through an INORGANIC MEMBRANE FILTER (Whatman^TM^ Anotop^TM^ 25/0.02, Cytiva), and was used as DNA template for polymerase chain reaction (PCR).

The β-tubulin gene (primers 237F–389R) of *H. akashiwo* H93616 (58) and the major capsid protein gene (*mcp*, 290F–429R) of HaV53 (21) (primers listed in **Table S2**) were amplified using TaKaRa Ex Taq DNA Polymerase (TaKaRa Bio), according to the manufacturer’s protocol. Reaction mixtures (details in **Table S4**) were briefly spun down and subjected to PCR using the Thermal Cycler Dice Touch (TaKaRa Bio) under the following conditions: 30 cycles of 3-step PCR (98°C for 10 s, 60°C for 30 s, 72°C for 15 s), followed by a final extension at 72°C for 3 min. PCR product amplification was confirmed by agarose gel electrophoresis using 2.5% agarose gels prepared with agarose S (NIPPON GENE) and stained with Gel Green Nucleic Acid Stain (Biotium) and imaged using a Gel Doc EZ Imager (Bio-Rad). In addition, PCR products were purified from the gel using the Wizard SV Gel and PCR Clean-Up System (Promega) according to the manufacturer’s protocols, and DNA concentrations were determined using the Qubit dsDNA HS Assay. Gene copy concentrations (copies/µL) were calculated based on the DNA concentration and amplicon length.

The standard DNA was serially diluted 10-fold in virus-free water for quantitative PCR. Reaction mixture composition and thermal cycling conditions were based on TaKaRa Bio’s protocol (details in **Table S5**) in 0.1 mL 8-strip PCR tubes (NIPPON GENETICS), using the Thermal Cycler Dice Real Time System III (TaKaRa) under the following conditions: 95°C for 30 s, 95°C for 5 s, 60°C for 30 s, 72°C for 15 s for 40 cycles, and dissociation 95°C for 15 s, 60°C for 30 s, 95°C for 15 s. The standard curve is generated using the second-derivative maximum method, based on the standard DNA concentration (copies/µL) and the cycle threshold (Ct) value. These curves were used to determine the copy number of β-tubulin and *mcp* gene in the extracted DNA samples.

### Investigation of *Aurantiochytrium* proliferation stimulated by lysis of virus-infected *H. akashiwo*, with host and virus quantification

To examine the impact of virus-infected *H. akashiwo* H93616 strain on the growth of *Aurantiochytrium* sp. NBRC102614, *H. akashiwo* H93616 was inoculated into 1.2 L of f/2-Si medium in a 2 L conical flask, until the cell density reached approximately 10^4^ cells/mL. For the HaV53 strain, 10 mL of HaV suspension was added to 90 mL of *H. akashiwo* H93616 culture medium in logarithmic growth phase in a 75 cm^2^ vented-cap type cell culture flask (ASONE) at the start of the light period. In terms of *Aurantiochytrium* sp. NBRC102614, this study followed and revised the previous method (56, 59), in which 20 mL of sterilized seawater was introduced to the d-GPY agar plate (containing glucose 2 g, peptone 1 g, yeast extract 0.5 g, and agar 15 g in 1 L artificial seawater) of *Aurantiochytrium* sp. NBRC102614 to wash off and collect cells containing zoospores.

After pre-culturing, 2.0 mL of *Aurantiochytrium* sp. NBRC102614 and its zoospores were introduced to the fresh flask. Subsequently, the *H. akashiwo* H93616 culture was transferred into six fresh flasks (178.2 mL per flask). For the Infection treatment, 19.8 mL of HaV53 virus was added, while the same volume of f/2-Si medium was added to the non-infection group, and as control, 198.0 mL of f/2-Si medium was added. These operations yielded the total volume to 200 mL in all treatments. Samples were collected from day 0 to day 8 for microscopic observation and DNA extraction. Specifically, samples were collected from the Infection, Non-infection, and Control group every 1 day, to conduct cell counting by microscopic observation of *Aurantiochytrium* sp. NBRC102614.

On the other hand, 2 mL of sample was collected from the infection and non-infection groups on days 0, 2, 4, 6, and 8 of culture in a 2 mL tube for DNA extraction. The mixture was centrifuged at 12,000 g × 5 min to pellet the cells. The supernatant was collected, and 2 mL of the culture medium was added. This procedure was repeated three times to pellet 6 mL of the culture medium. The pellet was suspended in 2 mL of f/2-Si medium, centrifuged at 12,000 g × 5 min, and the supernatant was discarded, with the procedure repeated thrice to remove extracellular HaV53. Simultaneously, 6 mL of the supernatant was collected from the infection group after centrifugation (12,000 g × 5 min). All the collected samples were stored at −80°C until DNA extraction. For cell density determination, 1 mL of the sample collected from each treatment was fixed with 1 mL of fixative (0.2 M sucrose, 5% glutaraldehyde in f/2-Si medium) and subsequently counted using a cell counting board (Wakenyaku). Significant differences between experimental time points were determined using the Mann–Whitney U test.

## CONCLUSION

This study demonstrates for the first time, using both environmental and culturing approaches, that heterotrophic Labyrinthulomycetes are key, previously underestimated, consumers of DOMderived from harmful algal blooms lysed by viral infection. In particular, this study revealed that Labyrinthulomycetes play a vital ecological role in shaping *Heterosigma* communities, as revealed by the amplicon analysis. Further investigation confirmed that large amounts of organic matter were likely released into the water column when algal cells were lysed by viral infection, enriching the pool of marine dissolved organic matter. Co-culturing *Aurantiochytrium* sp. with virus-infected *H. akashiwo* showed that algal cells were fully lysed by 3 days of infection, coinciding with a sharp increase in *Aurantiochytrium* cell density. After cell lysis, clear differences were observed in the growth of *Aurantiochytrium* sp. NBRC102614 indicated that the release of organic matter from lysed algal cells significantly promotes the growth of heterotrophic eukaryotes. These processes are likely to mediate efficient transfer of algal-derived organic matter to heterotrophic microbial consumers.

Taken together, our findings demonstrate that these mechanisms facilitate the transfer of virus-infected alga-derived organic matter from microalgae *H. akashiwo* to heterotrophic Labyrinthulomycetes and enhance their growth.

## ETHICAL COMPLIANCE

This research does not involve human subjects or animals.

## AUTHOR CONTRIBUTIONS

Takashi Yoshida, Ryoma Kamikawa, contributed to the experiment design, Miho Aoki and Shuhe Chen contributed to experiment implementation of the research, Shuhe Chen and Keishiro Sano contributed to the analysis of the results and to the writing of this manuscript. Takashi Yoshida, Ryoma Kamikawa, Keigo Yamamoto, and Yoshitake Takao conceived the original and supervised the project.

